# The Generalized Quantitative Model of antibody homeostasis: maintenance of global antibody equilibrium by effector functions

**DOI:** 10.1101/169276

**Authors:** József Prechl

**Affiliations:** Diagnostcum Zrt, Attila út 126, Budapest, Hungary MTA-ELTE Immunology Research Group, at Department of Immunology, ELTE, Pázmány P.s 1/c, Budapest, Hungary

**Keywords:** antibody, homeostasis, equilibrium, effector function, quantitative biology

## Abstract

The homeostasis of antibodies can be characterized as a balanced production, binding and elimination regulated by an interaction network, which controls B-cell development and selection. Recently we proposed a quantitative model to describe how the concentration and affinity of interacting partners generates a network.

Here we argue that this physical, quantitative approach can be extended for the interpretation of effector functions of antibodies. We define global antibody equilibrium as the zone of molar equivalence of free antibody and free antigen and immune complex concentrations and of dissociation constant of apparent affinity: [Ab]=[Ag]=[AbAg]=K_D_. This zone corresponds to the biologically relevant K_D_ range of reversible interactions. We show that thermodynamic and kinetic properties of antibody-antigen interactions correlate with immunological functions. The formation of stable, long-lived immune complexes correspond to a decrease of entropy and is a prerequisite for the generation of higher order complexes. As the energy of formation of complexes increases we observe a gradual shift from silent clearance to inflammatory reactions. These rules can also be applied to complement activation-related processes, linking innate and adaptive humoral responses. Affinity of the receptors mediating effector functions shows a corresponding range of affinities, allowing the continuous sampling of antibody-bound antigen over the complete range of concentrations. The generation of multivalent, multicomponent complexes triggers effector functions by cross-linking these receptors on effector cells with increasing enzymatic degradation potential.

Thus, antibody homeostasis is a thermodynamic system with complex network properties, nested into the host organism by proper immunoregulatory and effector pathways. Maintenance of global antibody equilibrium is achieved by innate qualitative signals modulating a quantitative adaptive immune system, which regulates molecular integrity of the host by tuning the degradation and recycling of molecules from silent removal to inflammatory elimination.

## Introduction

In a series of articles we present a generalized quantitative model for the homeostatic function of clonal humoral immune system, mapping antibody-antigen interactions in a virtual physical space. In our model we assumed that the essence of antibody (Ab) homeostasis is the maintenance of a regulated balance between the saturation of B-cell receptors (BCR) by antigen (Ag) and the saturation of antigen by antibodies, thereby controlling B-cell development and antigen fate in the body. Regulated antibody production drives the system of molecules towards a binding equilibrium, which characterizes the nature of immunological recognition. The formation of antibody-antigen complexes (AbAg) is followed by their removal from the body: in this third paper we describe how physical features of antibody binding to targets further expands and modulates the network of antibody interactions empowering the immune system with the control of the rate and mode of molecular degradation and reusal. We shall argue that our quantitative model supplements qualitative molecular approaches to immune function and provides a general physical framework with simple straightforward rules of operation.

## Global Antibody Equilibrium

We have defined humoral adaptive immunity as a system where qualitative signals of antigenic molecules adjust equilibrium dissociation constant K_D_ to be equal to free antigen concentration [Ag] via regulation of B-cell selection and differentiation ^1^. Quantitative signals drive B-cell proliferation and antibody production to adjust free antibody concentration [Ab]=K_D_ in order to achieve relevant Ag saturation ^2^. Therefore the system tends to reach K_D_=[Ag]=[Ab] at which point [AbAg] will also be equal to KD (Figure 1.) based on the general equation:

**Figure 1.**
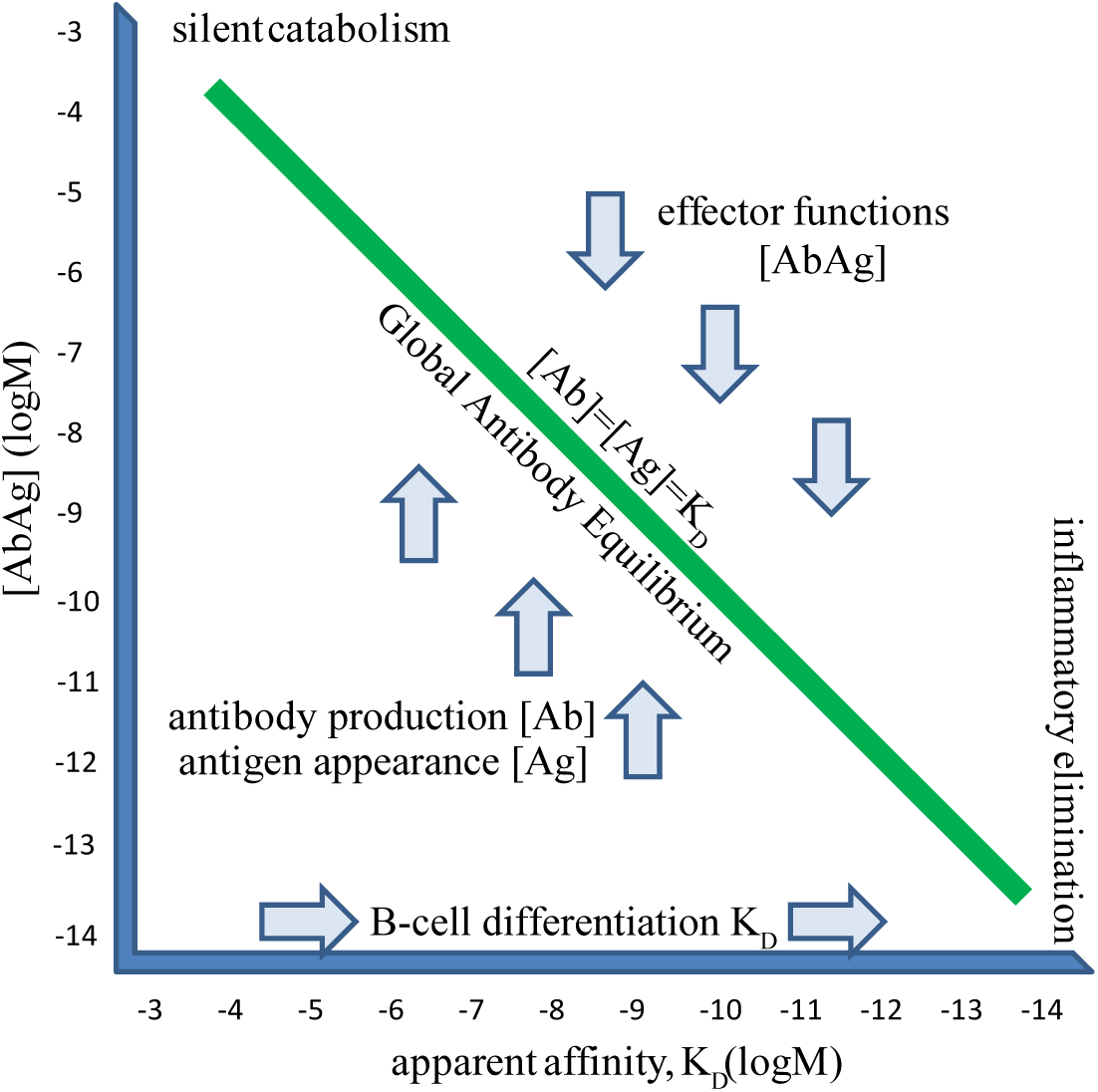
Equilibrium at equimolar antibody and antigen concentrations – the global antibody equilibrium. For each Kd value there is one special condition, when concentration of free and bound forms of Ab and Ag are all equal. For the complete range of immunologically relevant KDs and all interacting partners in the system the existence of a global equilibrium we call Global Antibody Equilibrium. The function of the humoral adaptive system is the maintenance of this equilibrium by adjusting Kd and [Ab] by B-cell proliferation and differentiation and [AbAg] by effector functions. This leads to th regulation of [Ag] according to its quality and quantity.

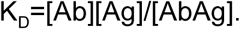

Thus, we can think of the clonal humoral immune system as one that controls [AbAg] by adjusting K_D_ and [Ab] according to changing [Ag] and the quality of Ag:

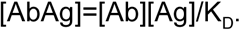

In turn this provides a means of controlling the rate and efficiency of removal of Ag from the host. Since the constant production of Ab and the continuous presence and appearance of Ag perpetually generates immune complexes, the only way of keeping [AbAg] constant is to remove it from the system. This is achieved by effector functions coupled to the humoral immune system - the rate, efficiency and quality of removal are key characteristics of and basically define immune responsiveness and responses.

## Physical aspects of immune complex formation

### Affinity and avidity

Antibody molecules bind to their targets via the paratope: a surface formed by the hypervariable peptide loops responsible for binding. The paratope contacts the epitope, the binding surface of the antigen, and the binding can be characterized by the affinity of the interaction. Multiple identical binding sites of antibodies can simultaneously bind to a multivalent target, carrying multiple identical epitopes. Recognition of a target with repetitive patterns is therefore promoted by placing several binding sites onto an antibody. Each site can bind to the paratope adding its affinity, the cumulative binding strength is called avidity. It is important to note that avidity is not a simple mathematical sum of affinities but rather a product of those. IgM, the first antibody produced by a B-cell, is pentameric or hexameric, with ten or twelve paratopes, respectively, allowing a significant shift in avidity upon multiple interactions (Fig.2). For the sake of consistency we will continue to use the expression apparent affinity as a general term for the characterization of the strength of molecular interactions, along with the use of K_D_.

**Figure 2.**
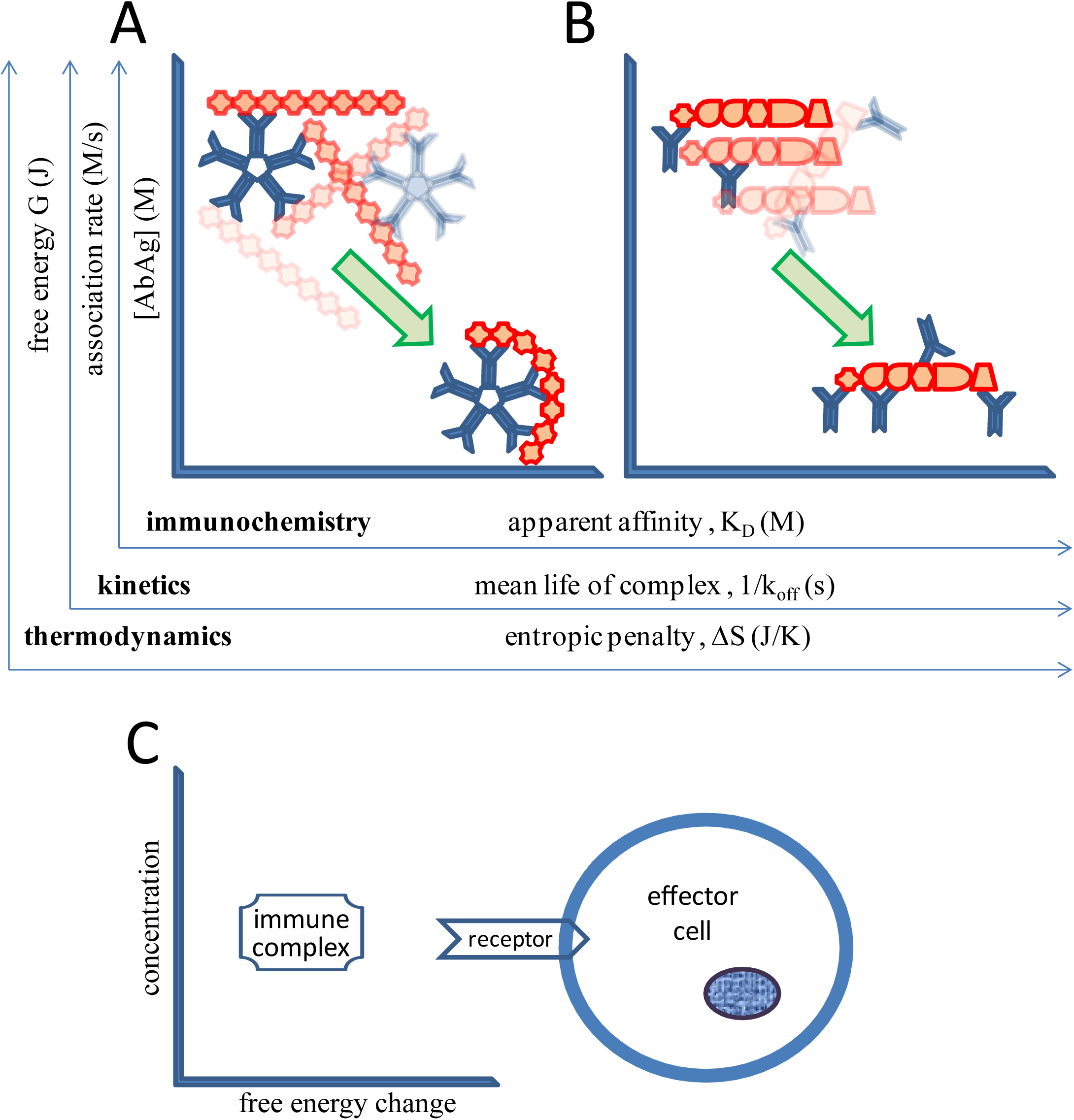
Interpretation and correlation of biophysical aspects of antibody-antigen interactions. Interactions of a single pentameric IgM (A) and several dimeric IgG (B) molecules are used as models. The same events can be interpreted from immunochemical, kinetic and thermodynamic aspects. Immune responses are charcterized by increased avidity – multivalent interactions between antibody and antigen result in higher apparent affinity; altered kinetics – the length of the interaction also increases with increased avidity; and different thermodynamic properties - high avidity interactions are characterized by higher free energy loss and greater decrease in entropy of the interacting molecules. Interaction of the complexes with cells is mediated by receptors of varying affinities, in turn cellular effects are determined by the extent of receptor cross-linkage and the identity of the effector cell.

In addition to this immunochemical interpretation K_D_= [Ab][Ag]/[AbAg], equilibrium can also be approached from the thermodynamic or from the kinetic aspect. Description of immunological effector functions requires an understanding of all of these approaches (Fig.2).

### Kinetics

The association rate constant k_on_ of the interacting molecules along with the dissociation rate constant k_off_ together describe where an equilibrium between free and bound forms is established, defined by the equation

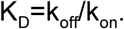

As suggested by the equation the same K_D_ could be the result of very fast and very slow event rates as long as their ratio remains constant. Since association rate constant has an upper limit due to diffusion limits ^3^ if we assume that k_on_ is similar in the complete range then the rate of association (rate=k_on_[Ab][Ag]) will be determined by the concentration of the interacting molecules. High concentrations of interacting Ab and Ag lead to frequent binding events (Fig.2A,B; kinetics). We can also assume that low affinity interactions are characterized by fast off rates, which at high concentrations of interactors translate to high frequency on-off reactions. High affinity interactions, on the other hand, are characterized by low dissociation rate constants, and low immune complex concentrations also result in low dissociation rates (rate=k_off_[AbAg]), meaning the generation of long-lasting complexes.

### Thermodynamics

Spontaneous reactions are accompanied by a decrease of free energy of the reactants ^4^. The relationship between affinity and the change in free energy is given by the equation:

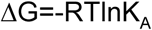

where ΔG stands for free energy change, R is molar gas constant (R = 8.314 J mol^−1^ K^−1^), T is thermodynamic temperature in Kelvins and K_A_ is the association rate constant. At constant temperature free energy change is thus proportional to affinity of the interaction. This change of free energy has enthalpic (ΔH) and entropic (ΔS) components

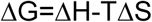

which can be used to describe the energy landscape of the antibody interaction network ^5^. In the next sections we will keep describing immune complex formation as a function of concentration and free energy change (Fig.2C), because we will be leaving the range of reversible interactions. One reason for that is the formation of higher order complexes with multiple sites of a single antibody binding (Fig.2A) or multiple antibodies binding (Fig.2B), the other reason being the appearance of covalent binding as we shall see in the next sections.

## FcR-mediated effector functions

Effector functions of antibodies are exerted via further interactions with cells. Neutralization might be regarded as an exemption, the neutralizing antibody binding to biologically important epitopes and interfering with their binding. However, even in this case complexes of the antigen and antibody need to be removed from the body at the end. Other effector functions, such as endocytosis, opsonin-mediated phagocytosis, frustrated phagocytosis, antibody-dependent cytotoxicity, complement-dependent cytotoxicity, all depend on interactions of immune complexes with effector cells. Primary mediators of antibody-cell interactions are the receptors binding the Fc region of immunoglobulins, FcR. These are characterized by their specificity for each antibody class and by their affinities. Here we considered FcR comprising one or more extracellular domains responsible for Fc binding that belong to the immunoglobulin superfamily (Fig.3). Reported affinities for FcμR ^6^ and FcαμR ^7^ are higher than shown here, a discrepancy possible caused by avidity effects and the use of transfected cells for affinity estimations. Nevertheless our theoretical estimations need experimental confirmation. Affinities of FcαRI ^8^, FcγRs ^9^ and FcεRI ^10, 11^ have been extensively studied, and while IgG subclasses introduce further level of complexity ^9,12^, a general gradual increase of affinity in this order is observed as shown in Figure 3.

**Figure 3.**
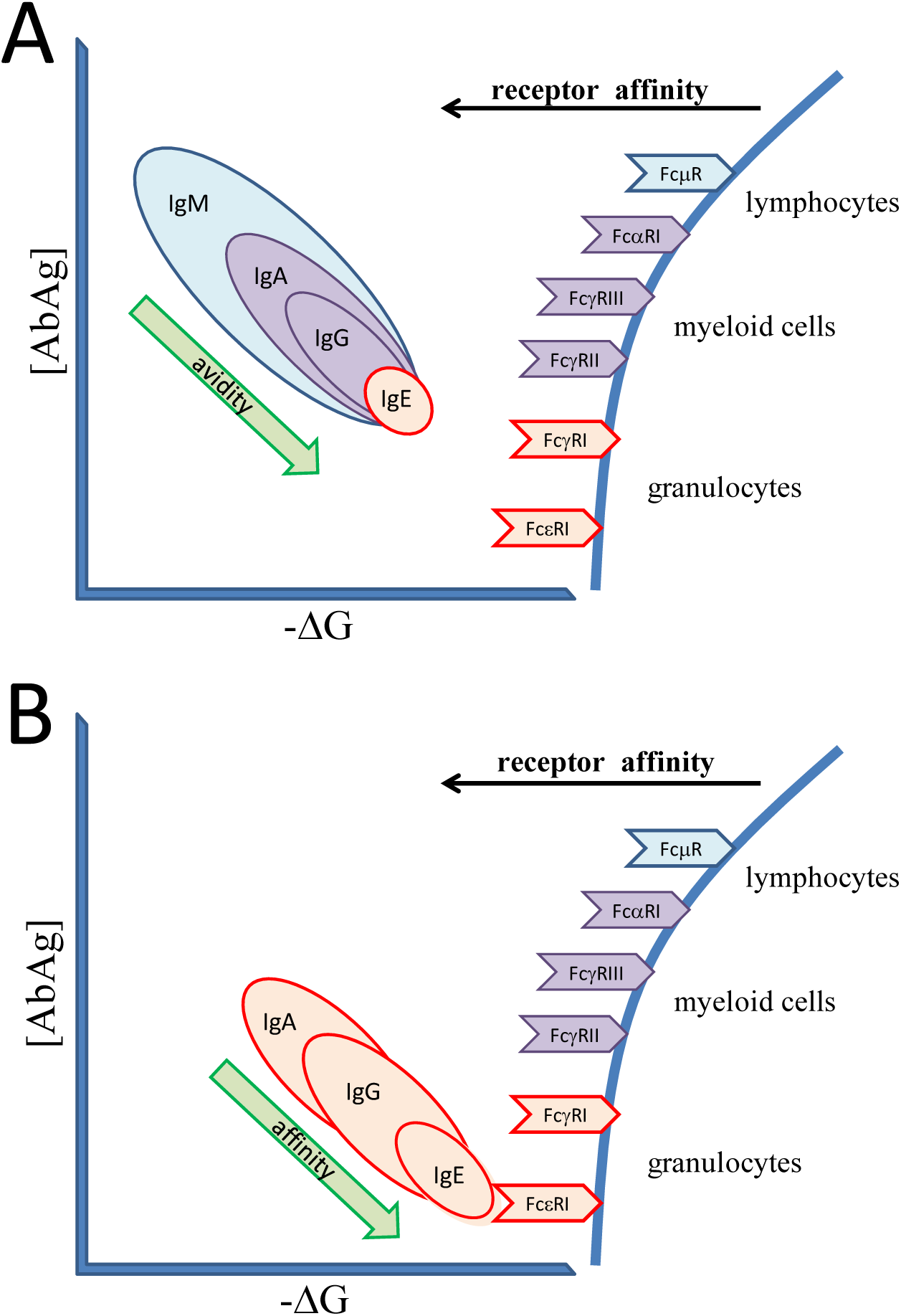
Effector functions of antibodies produced during TI and TD responses A, Thymus independent antibody responses operate by exploiting avidity effects: clones selected by specificity can increase their apparent affinity by multivalent binding to the target, if the target carries multiple identical epitopes. Hexameric IgM has twelve binding sites, dimeric IgA has four, while IgGs have two sites. Switching isotypes enhances cellular responses by binding to receptors with increasing affinity. B, Thymus dependent responses exploit affinity maturation besides isotype switching. Increased affinity to target is accompanied by the usage of isotypes with stronger effector functions. IgG3 is the most flexible of IgGs, while IgG4 may be functionally monovalent because of half-Ig swithcing. Affinity of IgE to FceRI is so high that cellular receptors are decorated with monomeric IgE. The presence of multiple high affinity antibodies on the target results in potent receptor cross-linkage, triggering robust cellular responses even via otherwise low-afinity receptors.

When considering the interactions of immune complexes with cells again we have to take into account not only the affinity but also the avidity of the interactions. Binding to multiple receptors by the same complex will cross-link receptors leading to enhanced signalling and more robust cellular responses. Monovalent engagement leads to homeostatic recycling in contrast to multivalent engagement, which can trigger cell activation via the FcR gamma chain ^8,13-15^.

Since we previously separately addressed thymus-independent (TI) and thymus-dependent (TD) antibody production ^2^, here we will also discuss effector functions using this categorization. We need to keep in mind however, that these events actually show an overlap and continuity when represented in our molecular interaction space (Fig.3A and B).

### TI responses

In the absence of T-cell derived co-stimulation B cells cannot initiate germinal center formation and cannot increase antibody affinity to target. TI responses therefore operate by taking advantage of avidity increase via multiple interactions, by changing antibody isotype and by adjusting antibody concentration (Fig.3A). IgM, the isotype displayed first by developing B cells is secreted in pentameric or hexameric form ^16^, the monomeric units covalently bound by interchain disulphide bridges or by the J chain ^17^. Polymeric IgM molecules have therefore ten to twelve antigen binding sites. Coupled with the flexibility of the molecule this results in multiple interactions with targets carrying repetitive epitopes. Such molecules are carbohydrates, glycoproteins, DNA and RNA, in addition to various microbial molecules ^18,19^. Free and bound IgM molecules bound by FcµR are endocytosed and shuttled to the lysosomes for degradation ^20^. Dimers of IgA (dIgA) form by J-chain incorporation, leading to tetravalent binding potential. Dimeric and monomeric IgA binds to CD89, also called FcαRI, which mediates phagocytosis and cellular activation ^21^. Priming of myeloid cells via IgG-mediated activation modulates FcαRI effects, a synergism potentially leading to inflammatory reactions ^22^. IgG2 is the characteristic isotype of responses against capsular bacteria ^12^. It has low affinity to most FcγRs, yet when bound to a multivalent target like polysaccharide we assume it can trigger substantial receptor cross-linkage. IgG1 is less characteristic for anti-polysaccharide responses and has higher affinity for FcγRs, potentially triggering cellular effector functions more efficiently. While IgE is mostly considered high affinity antibody, class switch to IgE antibodies independent of T-cell help has been shown in humans ^23^, suggesting its contribution to TI responses without affinity maturation and somatic hypermutation. In mice direct switching to IgE following TI stimulation was also observed, and low affinity IgE was shown to be protective against anaphylaxis ^24^. Effector functions triggered by TI antibody responses can thus range from silent uptake to robust phagocytosis, the response being modulated by immune complex avidity and quality and extent of receptor engagement (Fig.3A).

### TD responses

T cells provide co-stimuli for B cells, enabling affinity maturation and class-switching. TD responses are thus characterized by non-IgM isotypes - such as IgA, IgG and IgE - and increased affinity to target (Fig.3B). Higher affinity favours the generation of higher order complexes, since the longer duration of interactions allows several different antibodies to a single target with multiple epitopes. TD responses are characteristic for protein specific responses and proteins are usually not repetitive but show distinct highly variable epitopes. A polyclonal response targeting different epitopes will thus favour the generation of high avidity complexes with higher receptor-crosslinking potential. Granulocytes can arm themselves with antibodies binding to their high affinity receptors, FcαRI ^25^, FcγRI ^26,27^ and FcεRI, and degrade target antigen following receptor-crosslinkage, phagocytosis or exocytosis.

The majority of human IgE arises from class switched B cells already undergone somatic hypermutation ^28^. The affinity of allergen specific IgE in the nano- to picomolar K_D_ range has been found to correlate with cellular effects induced by antigen binding ^29^ and allergy phenotype has also been related to IgE affinity ^30^. Factors promoting receptor cross-linkage, such as increased avidity due to binding to multiple epitopes and relative content of specific to total IgE also enhanced degranulation of basophil granulocytes ^31^.

## Complement-mediated effector functions

The complement system is a phylogenetically ancient homeostatic system, predating the appearance of adaptive immunity. Accordingly, it can function independent of antibodies but it is also regulating and is regulated by the adaptive immune system ^32,33^. From our point of view an important aspect of complement-mediated immunity is that we can insert the whole system into the same framework as we did with antibodies (Fig.4). Recognition elements of the system are multivalent promiscuous binders ^34^, just as natural IgM. On the other hand, the complement system has numerous receptors and regulators, which are able to fine-tune effector functions ^35^. The classical pathway of complement can be initiated by C1q, a hexameric molecule, which binds antibodies as well as numerous self and non-self molecules ^36,37^. The lectin pathway is triggered by mannan-binding lectin, ficolins and other multimeric molecules ^38^. Less specific recognition is coupled to avidity effects by allowing multiple binding to repetitive patterns for all these molecules. While adaptive immunity needs time (days) to evolve and adapt to the antigen and tune the response, the complement system proceeds immediately to the next stages under appropriate conditions. Molecules bound to pattern recognition molecules can be swiftly and silently removed without further propagation of the cascade, the best example being the removal of apoptotic debris ^39^. Proteases attached to the recognition molecules can trigger proteolytic cleavage of further complement components, depending on the stability - as reflected by ΔG - of the complex. Covalent binding of C4b - a general feature of thioester-containing proteins ^40^ - further stabilizes the complex and provides ligand for other homeostatic receptors like CR3 and CR4 ^41^. In the absence of inhibitory and regulatory molecules, numerous complement C3 breakdown products are generated, leading to the formation of enzymatically active C3 and C5 convertases (Fig.4). Cleavage of C3 and C5 also generates fragments with anaphylatoxic properties acting on C3aR and C5aR, recruiting and activating leukocytes. These substances can trigger the degranulation of basophil, eosinophil granulocytes and mast cells, leading to massive systemic effects ^42^. Finally, a membrane-disrupting superstructure (Membrane Attack Complex) may form from the assembled complement components, lysing target cells.

**Figure 4.**
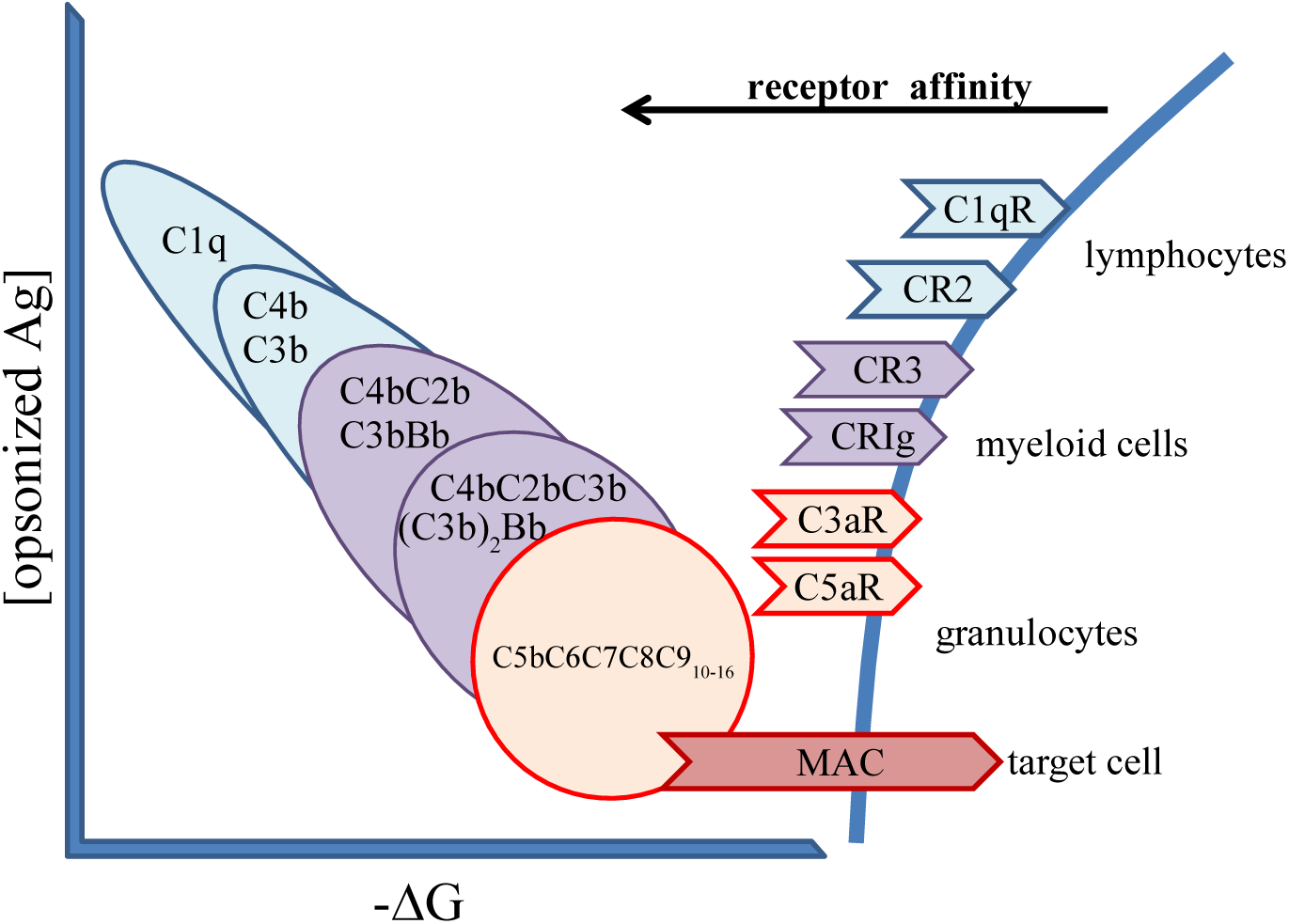
Effector functions of the complement system Innate homeostatic regulation by the complement system employ avidity effects, biochemical tagging and cell lytic effects. Initiators of the classical and lectin pathways are oligomeric molecules with mutiple binding sites. Attachment to the target by more than one arm triggers associated serine proteases and cleave complement proteins to initiate a cascade. Higher order complexes bound to the target cell can release chemotactic substances and form a membrane attack complex (MAC) with lytic function.

Thus, a sequence of events starting with avidity-mediated binding, assembly of structures with decreasing free energy and more robust cellular effects is observed, exactly as for antibodies, the difference being the lack of affinity and specificity maturation and a much faster flow of events (Fig.4).

## Interpretation of effector functions as degrees of antibody homeostasis

So far we have outlined a scheme wherein antibodies with varying binding affinities possess specific receptors with a corresponding range of affinities. It seems that varying serum concentrations of ligands (antibodies or immune complexes) can saturate their receptors to various extent, guaranteeing acontinuous homeostatic removal of antibodies for degradation. The concentration of serum IgM, and of IgA and IgG subclasses are all in the micromolar range ^43^ ensuring the binding to appropriate receptors with K_D_ in this range. As apparent K_D_ of both immune complex formation and receptor binding decreases the mean lifetime of immune complexes increases (Fig.2), leading to an increased chance of surface display and internalization of bound antigens. Both of these processes are important for the development of an immune response: cell surface presentation of intact antigen is critical for B-cell development, while internalization followed by enzymatic processing and MHC association is required for T-cell development and activation. As the number of recruited and engaged receptors increases the ensuing cellular responses are also tuned up. Receptor functions induced by signaling motifs in the polypeptide chains and by associated signaling molecules are effectively triggered when receptors are cross-linked by antigen-bound antibodies or multivalent antigens binding to surface-bound antibodies. This cross-linkage triggers robust cellular responses as determined by extent of cross-linkage, identity of the cell and coupling of the receptor to the cell.

FcμR are highly expressed by B-cells, allowing the internalization and degradation of circulating IgM molecules ^20^ as well as the presentation of bound antigen for other B-cells ^1^. An IgG receptor with inhibitory signaling motif, FcγRII is also expressed by B-cells, potentially allowing the surface presentation of antigen to other B cells, as has been observed for dendritic cells ^44^ Myeloid cells are more potent regarding degradatory effector functions; these cells also express FcγRIII and FcγRI, along with activatory isoform of FcγRII, and can carry out a wide range of regulatory and effector functions mediated by IgG isotypes ^12^. The receptor for IgA, FcαRI has been shown to mediate inhibitory or activatory effects depending on the extent of receptor crosslinking ^45,46^. Systemic level inflammatory reactions leading to anaphylaxis can be triggered by IgG or the anaphylatoxins C3a and C5a ^47^. The exceptionally high affinity of IgE to both its target and FcεRI can lead to highly stable antigen-antibody-receptor complexes and trigger degranulation of effector cells ^31^. Examination of the events following Ab binding to Ag using our proposed layers of immunological reactivity reveals that the further we leave behind immunological self the greater the energy of complex formation (Fig.5A). This is accompanied by a reduction in the number of potential microstates the involved molecules can take up, meaning the complexes becoming more and more rigid, characterized by decreased intrinsic entropy or in other words increased negentropy. Along with this increased negentropy we observe more and more pronounced molecular degradation by enzymatic processes, shifting endocytosis through phagocytosis to exocytosis. Accordingly, silent intracellular events aimed at recycling of molecular material are first replaced by enhanced local clearance activity and further on by systemic inflammatory effects induced by the release of various mediators (Fig.5B).

**Figure 5.**
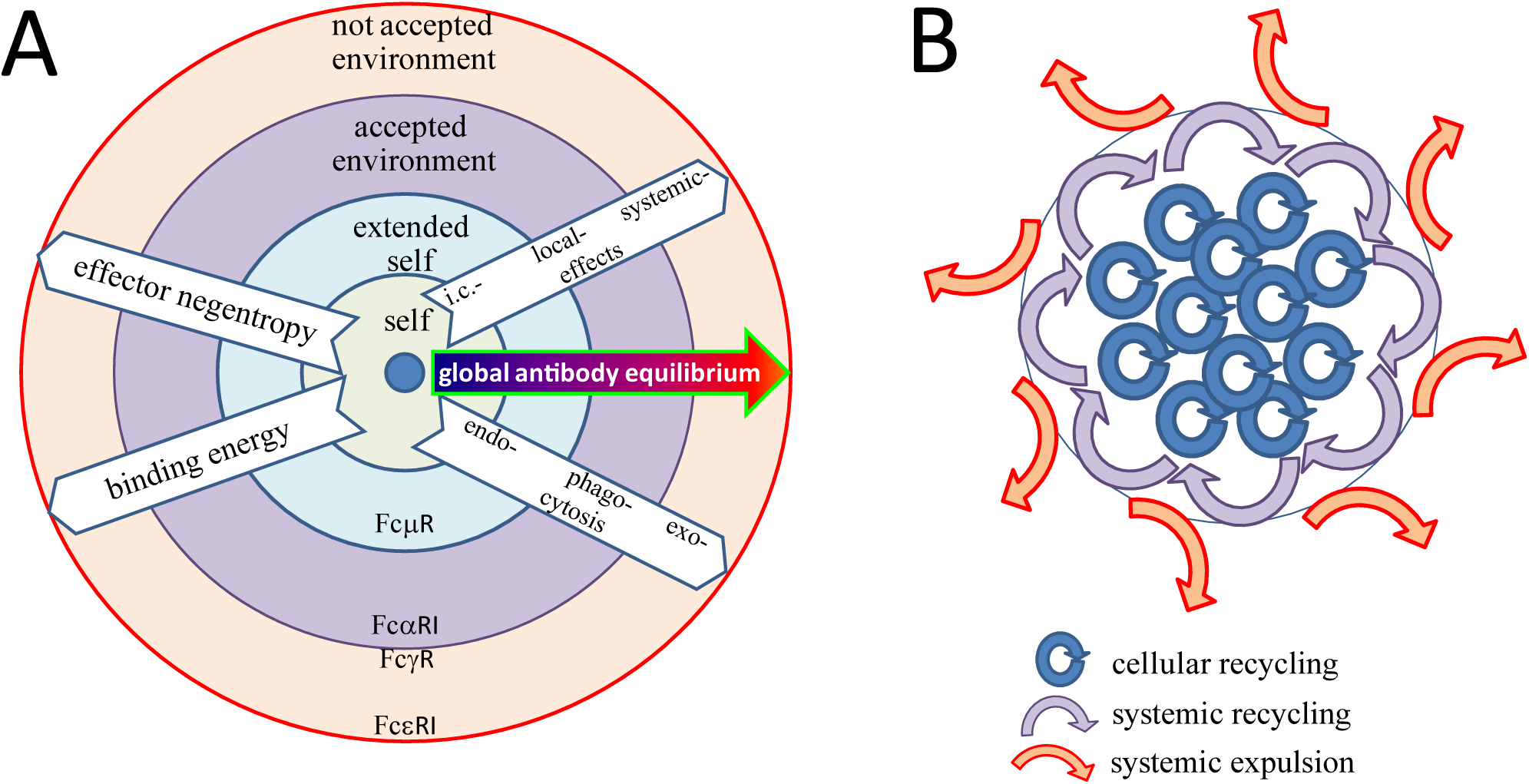
Antibody-mediated immune effector functions related to layers of immunological self. A, The generation of immune complexes with greater mass correlates with a greater loss of free energy and loss of entropy. As the entropy decreases the effector functions become more pronounced: intracellular lysosomal degradation shifts to cell activation and degranulation, with death of the effector and/or target cells. B, Homeostatic function of immune effector mechanisms represented as the flow of molecular material. Effector functions mediated via cellular receptors range from local recycling – silent uptake and recycling of immune complex components, through systemic degradation – lysosomal degradation, transport to organs, to expulsion – cellular destruction, inflammatory reactions. These events are positioned to reflect immunological layers of recognition, the circle representing our immunological border. i.c., intracellular

## Concluding thoughts

In two previous ^1,2^ and the present papers we have outlined a general scheme for the homeostasis of antibodies. The most important message of these papers is that humoral adaptive immunity can be viewed as a physical system responsible for the recycling of material. The biophysical role of the system is to achieve Global Antibody Equilibrium, a state when concentrations of free antibody, free antigen and bound antibody-antigen complexes are all equal to each other and also to the K_D_ of the interaction. The immunological role is to use qualitative signals of innate immunity for setting the apparent K_D_ for each interaction, thereby determining fate and life-time of the target molecules. The cell biological role is to remove antibodies and bound antigen, and degrade these molecules with appropriate efficiency. The whole system maintains body integrity by setting and stabilizing molecule concentrations over several orders of magnitude (Fig.1), depending on the quality and quantity of the molecules. This range of concentrations, affinities, binding energies is placed exactly where reversible macromolecular interactions occur. Once molecules exit this scale of interactions irreversible reactions take place: molecules are degraded in enzymatic compartments of various sizes, allowing the reuse of organic material.

After all, we think that the word immunity is actually somewhat misleading for describing the function of the adaptive immune system, since it comes from the latin “exempt, protected”. The adaptive humoral immune system, owing to its ability of cellular evolution via genetic diversification, is an ingenious physiological molecular balance-keeper of multicellular organisms, a system for the global regulation of all molecules, a continuum of recognition, complex formation and complex removal being present from silent self-maintenance to loud non-self expulsion. It is the qualitative signals of innate immune system, which superimposed on this physiological system render it suitable for protective purposes. It seems that maintenance of self and expulsion of non-self is the very same process, these entities being at the ends of a continuum. Indeed it seems intuitive that molecules that persist for long constitute the organism, while those that do not survive cannot be considered a part of it. Control of the quantity and life-time of molecules and molecular assemblies practically defines an organism - this is achieved by the humoral clonal immune system.

## Acknowledgments

I wish to thank my students and colleagues for their devoted experimental work and scientific discussions leading to the preparation of this manuscript.

## Bibliography

1 Prechl J. A generalized quantitative antibody homeostasis model: regulation of B-cell development by BCR saturation and novel insights into bone marrow function. Clin Transl Immunology 2017; 6: e130.

2 Prechl J. A generalized quantitative antibody homeostasis model: antigen saturation, natural antibodies and a quantitative antibody network. Clin Transl Immunology 2017; 6: e131.

3 Corzo J. Time, the forgotten dimension of ligand binding teaching. Biochem Mol Biol Educ 2006; 34: 413–6.

4 Smith RD, Engdahl AL, Dunbar JB, Carlson HA. Biophysical limits of protein-ligand binding. J Chem Inf Model 2012; 52: 2098–106.

5 Prechl J. Thermodynamic projection of the antibody interaction network: the fountain energy landscape of binding. BioRxiv 2017; published online April 5. DOI: 10.1101/124503.

6 Kubagawa H, Oka S, Kubagawa Y, et al. Identity of the elusive IgM Fc receptor (FcmuR) in humans. J Exp Med 2009; 206: 2779–93.

7 Shibuya A, Sakamoto N, Shimizu Y, et al. Fc alpha/mu receptor mediates endocytosis of IgM-coated microbes. Nat Immunol 2000; 1: 441–6.

8 Wines BD, Sardjono CT, Trist HH, Lay CS, Hogarth PM. The interaction of Fc alpha RI with IgA and its implications for ligand binding by immunoreceptors of the leukocyte receptor cluster. J Immunol 2001; 166: 1781–9.

9 Bruhns P, Iannascoli B, England P, et al. Specificity and affinity of human Fcgamma receptors and their polymorphic variants for human IgG subclasses. Blood 2009; 113: 3716–25.

10 Miller L, Blank U, Metzger H, Kinet JP. Expression of high-affinity binding of human immunoglobulin E by transfected cells. Science 1989; 244: 334–7.

11 Wan T, Beavil RL, Fabiane SM, et al. The crystal structure of IgE Fc reveals an asymmetrically bent conformation. Nat Immunol 2002; 3: 681–6.

12 Vidarsson G, Dekkers G, Rispens T. IgG subclasses and allotypes: from structure to effector functions. Front Immunol 2014; 5: 520.

13 van Egmond M, van Spriel AB, Vermeulen H, Huls G, van Garderen E, van de Winkel JG. Enhancement of polymorphonuclear cell-mediated tumor cell killing on simultaneous engagement of fcgammaRI (CD64) and fcalphaRI (CD89). Cancer Res 2001; 61: 4055–60.

14 van Vugt MJ, Heijnen AF, Capel PJ, et al. FcR gamma-chain is essential for both surface expression and function of human Fc gamma RI (CD64) in vivo. Blood 1996; 87: 3593–9.

15 Dombrowicz D, Flamand V, Miyajima I, Ravetch JV, Galli SJ, Kinet JP. Absence of Fc epsilonRI alpha chain results in upregulation of Fc gammaRIII-dependent mast cell degranulation and anaphylaxis. Evidence of competition between Fc epsilonRI and Fc gammaRIII for limiting amounts of FcR beta and gamma chains. J Clin Invest 1997; 99: 915–25.

16 Randall TD, King LB, Corley RB. The biological effects of IgM hexamer formation. Eur J Immunol 1990;20:1971–9.

17 Randall TD, Parkhouse RM, Corley RB. J chain synthesis and secretion of hexameric IgM is differentially regulated by lipopolysaccharide and interleukin 5. Proc Natl Acad Sci U S A 1992; 89: 962–6.

18 Lobo PI. Role of Natural Autoantibodies and Natural IgM Anti-Leucocyte Autoantibodies in Health and Disease. Front Immunol 2016; 7: 198.

19 Muthana SM, Gildersleeve JC. Factors Affecting Anti-Glycan IgG and IgM Repertoires in Human Serum. Sci Rep 2016; 6: 19509.

20 Lloyd KA, Wang J, Urban BC, Czajkowsky DM, Pleass RJ. Glycan-independent binding and internalization of human IgM to FCMR, its cognate cellular receptor. Sci Rep 2017; 7: 42989.

21 Morton HC, Brandtzaeg P. CD89: the human myeloid IgA Fc receptor. Arch Immunol Ther Exp (Warsz) 2001; 49: 217–29.

22 Kecse-Nagy C, Szittner Z, Papp K, et al. Characterization of NF-*κ* B Reporter U937 Cells and Their Application for the Detection of Inflammatory Immune-Complexes. PLoS ONE 2016; 11: e0156328.

23 Berkowska MA, Heeringa JJ, Hajdarbegovic E, et al. Human IgE(+) B cells are derived from T cell-dependent and T cell-independent pathways. J Allergy Clin Immunol 2014; 134: 688–697.e6.

24 Xiong H, Dolpady J, Wabl M, Curotto de Lafaille MA, Lafaille JJ. Sequential class switching is required for the generation of high affinity IgE antibodies. J Exp Med 2012; 209: 353–64.

25 Heineke MH, van Egmond M. Immunoglobulin A: magic bullet or Trojan horse? Eur J Clin Invest 2017; 47: 184–92.

26 Ioan-Facsinay A, de Kimpe SJ, Hellwig SMM, et al. FcgammaRI (CD64) contributes substantially to severity of arthritis, hypersensitivity responses, and protection from bacterial infection. Immunity 2002; 16: 391–402.

27 Swisher JFA, Feldman GM. The many faces of Fc *γ* RI: implications for therapeutic antibody function. Immunol Rev 2015; 268: 160–74.

28 Looney TJ, Lee J-Y, Roskin KM, et al. Human B-cell isotype switching origins of IgE. J Allergy Clin Immunol 2016; 137: 579–586.e7.

29 Mita H, Yasueda H, Akiyama K. Affinity of IgE antibody to antigen influences allergen-induced histamine release. Clin Exp Allergy 2000; 30: 1583–9.

30 Wang J, Lin J, Bardina L, et al. Correlation of IgE/IgG4 milk epitopes and affinity of milk-specific IgE antibodies with different phenotypes of clinical milk allergy. J Allergy Clin Immunol 2010; 125: 695–702, 702.e1.

31 Christensen LH, Holm J, Lund G, Riise E, Lund K. Several distinct properties of the IgE repertoire determine effector cell degranulation in response to allergen challenge. J Allergy Clin Immunol 2008; 122: 298–304.

32 Erdei A, Prechl J, Isaák A, Molnár E. Regulation of B-cell activation by complement receptors CD21 and CD35. Curr Pharm Des 2003; 9: 1849–60.

33 Sörman A, Zhang L, Ding Z, Heyman B. How antibodies use complement to regulate antibody responses. Mol Immunol 2014; 61: 79–88.

34 Degn SE, Thiel S. Humoral pattern recognition and the complement system. Scand J Immunol 2013; 78: 181–93.

35 Zipfel PF, Skerka C. Complement regulators and inhibitory proteins. Nat Rev Immunol 2009; 9: 729–40.

36 Gaboriaud C, Frachet P, Thielens NM, Arlaud GJ. The human c1q globular domain: structure and recognition of non-immune self ligands. Front Immunol 2011; 2: 92.

37 Nayak A, Pednekar L, Reid KBM, Kishore U. Complement and non-complement activating functions of C1q: a prototypical innate immune molecule. Innate Immun 2012; 18: 350–63.

38 Garred P, Genster N, Pilely K, et al. A journey through the lectin pathway of complement-MBL and beyond. Immunol Rev 2016; 274: 74–97.

39 Benoit ME, Clarke EV, Morgado P, Fraser DA, Tenner AJ. Complement protein C1q directs macrophage polarization and limits inflammasome activity during the uptake of apoptotic cells. J Immunol 2012; 188: 5682–93.

40 Shokal U, Eleftherianos I. Evolution and Function of Thioester-Containing Proteins and the Complement System in the Innate Immune Response. Front Immunol 2017; 8: 759.

41 Lukácsi S, Nagy-Baló, Z, Erdei A, Sándor N, Bajtay Z. The role of CR3 (CD11b/CD18) and CR4 (CD11c/CD18) in complement-mediated phagocytosis and podosome formation by human phagocytes. Immunol Lett 2017; published online May 26. DOI: 10.1016/j.imlet.2017.05.014.

42 Bosmann M, Ward PA. Role of C3, C5 and anaphylatoxin receptors in acute lung injury and in sepsis. Adv Exp Med Biol 2012; 946: 147–59.

43 Prechl J, Papp K, Erdei A. Antigen microarrays: descriptive chemistry or functional immunomics? Trends Immunol 2010; 31: 133–7.

44 Dubois B, Caux C. Critical role of ITIM-bearing FcgammaR on DCs in the capture and presentation of native antigen to B cells. Immunity 2005; 23: 463–4.

45 Ben Mkaddem S, Rossato E, Heming N, Monteiro RC. Anti-inflammatory role of the IgA Fc receptor (CD89): from autoimmunity to therapeutic perspectives. Autoimmun Rev 2013; 12: 666–9.

46 Bakema JE, van Egmond M. The human immunoglobulin A Fc receptor Fc *α* RI: a multifaceted regulator of mucosal immunity. Mucosal Immunol 2011; 4: 612–24.

47 Finkelman FD, Khodoun MV, Strait R. Human IgE-independent systemic anaphylaxis. J Allergy Clin Immunol 2016; 137: 1674–80.

